# Single molecule imaging reveals the concerted release of myosin from regulated thin filaments

**DOI:** 10.1101/357202

**Authors:** A. V. Inchingolo, M. Mihailescu, D. Hongsheng, N. M. Kad

**Keywords:** Cooperativity, muscle, actin, fluorescence imaging, regulation, calcium

## Abstract

Regulated thin filaments (RTFs) tightly control striated muscle contraction through calcium binding to troponin, which in turn shifts the position of tropomyosin on actin to expose myosin binding sites. The binding of the first myosin holds tropomyosin in a position such that more myosin binding sites on actin are available, resulting in cooperative activation. Troponin and tropomyosin also act to turn off the thin filament; however, this is antagonized by the high local concentration of myosin, questioning how the thin filament relaxes. To provide molecular details of deactivation we use the RTF tightrope assay, in which single RTFs are suspended between pedestals above a microscope coverslip surface. Single molecule imaging of GFP tagged myosin-S1 (S1-GFP) is used to follow the activation of RTF tightropes. In sub-maximal activation conditions, S1-GFP molecules bind forming metastable clusters, from which release and rebinding of S1-GFP leads to prolonged activation in these regions. Because the RTFs are not fully active we are able to directly observe deactivation in real time. Using a Reversible Jump Markov Chain Monte Carlo model we are able to dynamically assess the fate of active regions. This analysis reveals that myosin binding occurs in a stochastic stepwise fashion; however, an unexpectedly large probability of multiple simultaneous detachments is observed. This suggests that deactivation of the thin filament is a coordinated, active process.

## Introduction

Striated muscle contraction is mediated by the interaction of myosin II with actin and is governed by the calcium concentration within the myocyte. In the sarcomere, myosin II is assembled into thick filaments, which cyclically bind to the actin-containing thin filaments in the presence of ATP to generate force. However, access to actin is modulated by the control proteins troponin (Tn) and tropomyosin (Tm). Tn is a complex of three proteins that interact with both actin and the 40 nm long filamentous protein Tm. Calcium binding triggers a conformational change in Tn that permits Tm to slide across actin (1). This movement exposes sites on actin for myosin to bind, which in turn leads to the further exposure of myosin binding sites on actin (2). Together, these interactions result in cooperative activation of the thin filament and the generation of force. A further level of force control is provided by the thick filament, which responds to environmental cues to alter the amount of available myosin (3, 4). How muscle deactivates is as important as its activation, and defects in this process have been implicated in cardiac disease (5).

Activation of the thin filament is hypothesised to be modulated by the accessibility of myosin binding sites on actin within the context of a three-state mechanism (6). In the first of these states (blocked) tropomyosin sterically impedes myosin from binding to actin. Upon calcium binding to troponin, tropomyosin shifts across actin, partially uncovering myosin binding sites allowing a weak interaction between myosin and actin (closed state). Finally, in the open state, myosin binds strongly to actin and can generate force. Due to the stiffness of Tm, its movement to the open state also uncovers further myosin binding sites on actin, leading to cooperative activation (7). Since actin is a long filamentous molecule, there is no clear structural definition of the cooperative unit size. Instead this size is defined functionally in terms of the amount of actin exposed for myosin to bind (8–10). These and other studies (11, 12) suggest that a single myosin binding to a thin filament is capable of holding Tm open for the association of up to 7-14 further myosins. In reverse, this process is less clear. As the calcium concentration drops, how does Tm return to the blocked state in the presence of myosins at a very high local concentration (> 0.15 mM (13))?

Using a single molecule approach to measure the binding and release of fluorescent myosin molecules to thin filaments in sub-maximal calcium conditions, we are able to shed light on the process of deactivation. In this recently developed assay (9), single thin filaments are suspended between surface immobilized beads to enable access of myosin to actin unimpeded by a surface. With sufficient myosin, at the mid-point of calcium-induced activation (pCa_50_), a metastable activation process is observed. Myosin binds to regions of actin for long enough to permit additional myosins to bind in close proximity, without favouring complete propagation of activation as seen at saturating calcium (9). Here, we temporally resolve the interactions of fluorescently tagged myosin S1 with the thin filament. Changes in fluorescence intensity within active regions correspond to the release or binding of myosins. By following these events we can statistically reconstruct the fate of active regions. To enable an assumption-free model of the attachment and detachment probabilities for myosin, we modelled the observed binding and release of individual myosins using a Gaussian mixtures model, analysed via a Reversible Jump Markov Chain Monte Carlo approach. We found that, as expected, myosin binds to actin stochastically and forms clusters. However, a much larger than expected probability for simultaneous release of all myosins within an active cluster was observed in the data. In the metastable conditions studied, this was the most probable outcome for an active region. This highly elevated collapse probability suggests a concerted mechanism of deactivation (relaxation), and explains the ability of muscle to relax in conditions that would be expected to still permit myosin binding.

## Materials and Methods

### Proteins

Myosin and actin were prepared from chicken pectoralis following the protocol of Pardee and Spudich (14). Both human Tropomyosin and human Troponin were bacterially expressed in *E. coli.* Tropomyosin with an N-terminal Ala-Ser modification (to mimic acetylation (15)) was purified as follows: cell lysate was heated to 80°C for 10 min before centrifuging at 17300 g for 30 min at 4°C. The supernatant was aspirated and the pH lowered to 4.8, causing Tm to precipitate. After centrifugation at 3000 g for 10 min at 4°C, the pellet was resuspended in 5 mM potassium phosphate pH 7, 100 mM NaCl, 5 mM MgCl_2_ before further purification using a HiTrap Q HP column (GE Healthcare). This procedure was repeated once more and the pure protein stored in 20 mM Tris pH 7, 100 mM KCl, 5 mM MgCl_2_ at −80 °C.

The troponin complex was purified according to Janco and co-workers (16). Briefly, cell pellets for each of TnI, TnC and TnT were resuspended in 6 M Urea, 25 mM Tris pH 7, 200 mM NaCl, 1 mM EDTA, 20% sucrose and 0.1% Triton X100 and sonicated. Following centrifugation at 17300 g for 30 min at 4°C, the supernatants were combined and subsequently dialysed into 2 M Urea, 10 mM Imidazole pH 7, 1 M KCl and 1 mM DTT for 5 hours at 4°C. To allow the proteins to refold, the urea was removed by a second dialysis into 10 mM Imidazole pH 7, 0.75 M KCl, 1 mM DTT overnight at 4°C before a final dialysis into 10 mM Imidazole pH 7, 0.5 M KCl, 1 mM DTT for 5 hours at 4°C. The samples were then centrifuged at 23000 g for 10 minutes at 4°C and the protein in the supernatant precipitated by adding 30% ammonium sulphate (NH_4_)_2_SO_4_ and gently stirring at 4°C for one hour. After centrifugation at 8000 g for 30 minutes at 4°C, the sample was precipitated on ice for 30 minutes by bringing the ammonium sulphate concentration to 50%. The solution was centrifuged at 23000 g for 10 minutes at 4°C, and the pellet resuspended in 10 mM Imidazole pH 7, 200 mM NaCl, 100 μM CaCl_2_, 1 mM DTT before an overnight dialysis in the same buffer to remove the (NH_4_)_2_SO_4_. After dialysis, the sample was spin concentrated in a 10 kDa MWCO column (Amicon Ultra) at 2360 g for 45 minutes and further purified using a Sephacryl S-300 column (GE Healthcare). Finally, the protein sample was stored in the same buffer, with the addition of 3% sucrose, at −80°C. Reconstituted regulated thin filaments (RTFs), i.e. actin fully decorated with Tm and Tn, were obtained through an overnight incubation at 4°C in reconstitution buffer as described previously (17), with an Actin:Tm:Tn ratio of 2:0.5:0.25.

S1-GFP was prepared as described in (9). The S1 region of the myosin heavy chain was digested using 1 mg/ml papain (Sigma-Aldrich) in 5 mM cysteine pH 6 and 2 mM EDTA for 15 min at room temperature (18). The reaction was stopped with 5 mM iodoacetic acid and purified using a DEAE FF column (GE Healthcare). Regulatory light chain (RLC) C-terminally fused with a green fluorescent protein and a His-tag (RLC-GFP-6xHis) was recombinantly expressed in *E. coli* and purified using a Ni-NTA column. The purified construct was then exchanged for myosin S1 RLC (1:3 S1:RLC-GFP-6xHis ratio) in 50 mM potassium phosphate buffer pH 7, 600 mM KCl, 10 mM EDTA, 2.5 mM EGTA, 2 mM ATP for 1 hour at 30°C. The reaction was stopped with 15 mM MgCl_2_ and the construct further purified using a Sephacryl S-200 and a HisTrap HP (GE Healthcare). Purified proteins were then stored in 50% glycerol and 3% sucrose at −20 °C.

### Creating and imaging thin filament tightropes

RTFs were suspended between 5 μm silica beads (MicroSil™ microspheres, Bang Laboratories) coated in poly-L-lysine adhered to coverslip of a microfluidic flow cell (Figure 1a), as detailed in (19). These RTF tightropes were illuminated using a continuous wave variable power 20 mW 488 nm DPSS laser (JDSU), focused off-centre at the back-focal plane of a 100x objective (1.45 NA) to generate an obliquely angled field (20–22), custom-built into an Olympus IX50 microscope. Fluorescence was measured using a Hamamatsu OrcaFlash 4.0 camera, observing the interaction of S1-GFP with RTFs at pCa 6, 0.1 μM ATP and 5 nM S1-GFP. To reduce background noise from surface adhered S1-GFPs the surface of the flow-cell was photobleached for 1 min at 20 mW prior to recording data. Movies were recorded at 3.3 frames/second for 2 minutes exciting the GFP with 5 mW laser power, resulting in fluorescent spots where the S1-GFP co-localises with a RTF (Figure 1b). To further improve the signal/noise ratio the movies were binned 2×2, resulting in a pixel size of 126.4 nm.

**Figure 1. Imaging individual GFP tagged myosins interacting with regulated thin filaments.** ***a***) Regulated thin filaments (RTFs) are suspended between surface immobilized beads (top) using a microfluidic chamber (bottom). The silica beads are functionalised with poly-L-lysine that adheres them to the passivated (mPEG coated) coverslip surface and to the RTFs. Illumination is achieved at an oblique angle to reduce background and fluorescence detection occurs through the objective lens. b) Top-down view of S1-GFP molecules bound to an RTF suspended between silica beads (dashed circles). Each pixel is 126.4 × 126.4 nm.

### Data treatment and analysis

Movies of S1-GFP binding to and releasing from thin filament tightropes were converted into kymographs along the contour of actin using ImageJ. Kymographs provide a simple means of visualising the time domain of activation; each interaction of myosin with the RTFs appears as a horizontal line. The dosage of laser illumination, which is a product of intensity and duration, did not change interaction lifetimes of individual myosins indicating that photobleaching was not affecting this study in these conditions (9). Any systematic asymmetries in background noise were removed from the kymographs using ImageJ rolling ball background subtraction (radius = 300 pixels).

Extraction of fluorescence intensities from the kymographs was performed using an automated Matlab routine that fits each vertical slice (corresponding to a frame of the movie) to an autonomously-determined number of Gaussian distributions. The binding and release of myosin during periods of metastable activity (see results) leads to a shift in the mean position of the active region over time. Therefore, it was necessary to manually assign fluorescence peaks to active regions. Such idealized datasets could then be analysed using a Gaussian mixtures model via a Reversible Jump Markov Chain Monte Carlo (RJMCMC) approach, using custom-written software in R that utilizes the package miscF (23) and coda (24). Based on the results of the mixtures model analysis, we can further extract the transition matrix information for the stochastic process of myosin’s attachment and detachment. More details of the modelling can be found in the supplementary information.

## Results

### Generation and qualitative assessment of metastable acto-myosin interactions

Single molecule imaging of S1-GFP loading and release from RTFs offers a direct spatial and temporal view of how activation occurs. In our previous work (9), we developed a new approach to follow this process on RTF tightropes. These structures consist of individual RTFs suspended between beads adhered to a microscope coverslip surface (Figure 1a). The RTFs are constructed using microfluidics which are also used to add the assay components. In the presence of S1-GFP, we are able to detect binding to the RTF using fluorescence detection of the GFP; only those molecules binding to the tightrope are stationary long enough to register as a signal (Figure 1b). Previously, S1-GFP was seen to bind in clusters; the occupancy of these active regions was dependent on the concentration of myosin, ATP and calcium in the steady-state. Here, using the same experimental approach, we measure the dynamics of S1-GFP binding to and releasing from active regions. To increase the occurrence of formation and loss of active regions on the RTF we modulated [ATP] to prolong attachment while lowering the calcium concentration to near the pCa_50_ (mid-point of activation). This sub-maximal activation of the RTF prevents it from being fully turned on, leading to the formation of metastable active regions (Figure 2a). Examination of these metastable active regions indicates that clusters of S1-GFP form that can move laterally along the thin filament (Figure 2a) with no directional bias. This indicates that the binding/release of S1-GFP occurs stochastically with equal probability at both ends of the cluster (9). Since the intensity within the active region is directly related to the number of myosins present we can time-resolve the loading and release of myosin. In active regions we observe that binding occurs predominantly stepwise (Figure 2b), whereas detachment occurs both stepwise or through simultaneous detachment of multiple myosin molecules. This qualitative assessment of the data prompted a more robust examination using an unbiased statistical approach to determine the process of detachment.

**Figure 2.**
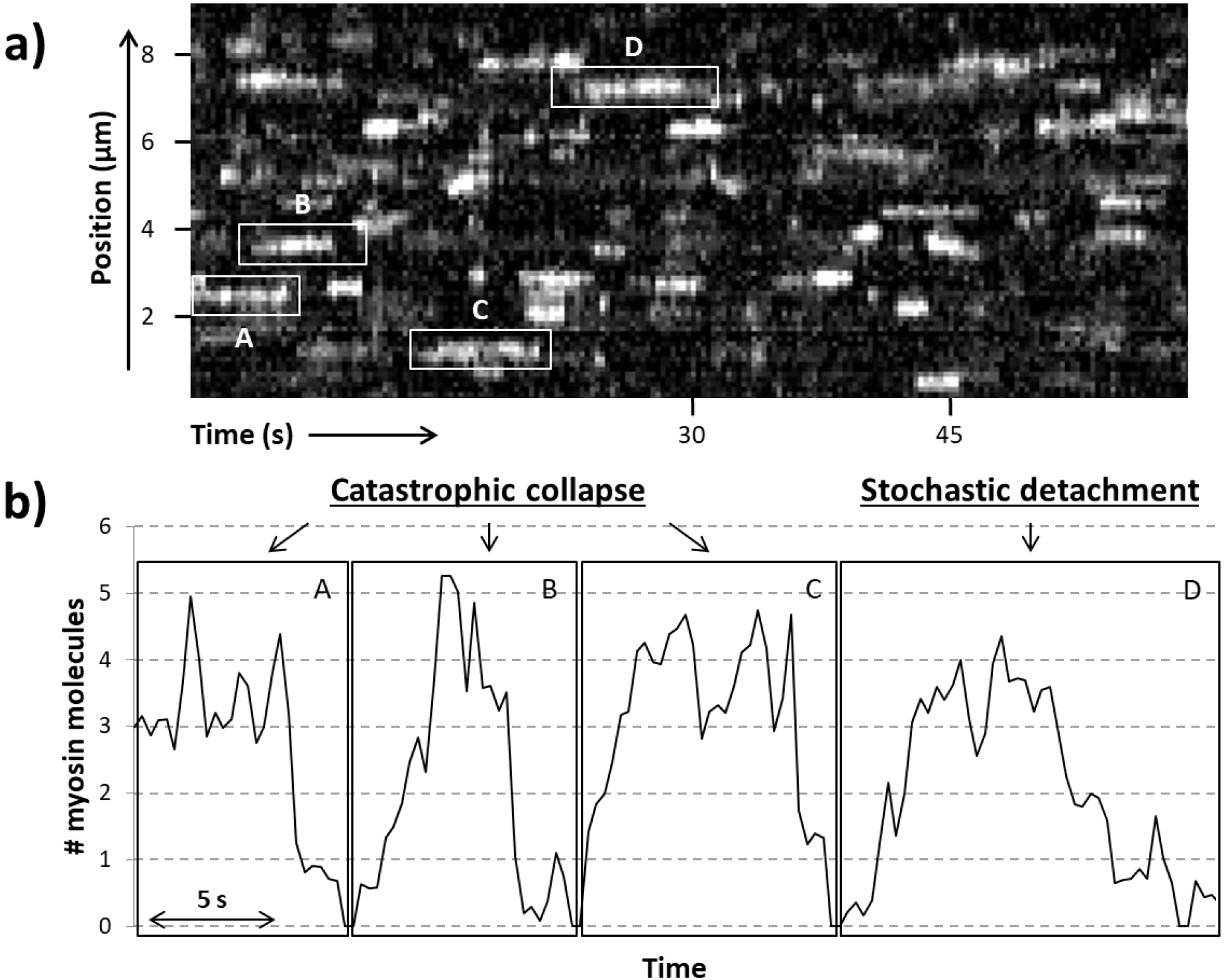
Active regions on an RTF tightrope. ***a***) A representative kymograph of an RTF in sub-maximal activating conditions, with some of the locally active regions (highlighted in boxes) labelled A to D. ***b***) The number of S1-GFP molecules bound in the highlighted active regions is shown. Sudden collapse of the active region is seen in particular for areas A, B and C, whereas region D shows both stepwise attachments and detachments. Data were obtained for 5 nM S1-GFP at pCa 6 and 0.1 μM ATP.

### Defining transition thresholds from RJMCMC analysis

To quantify the binding and release of S1-GFP within active regions we applied a Markov Chain statistical approach. The first step in our analysis was to identify and assign a peak intensity for each point in time for every active region in a kymograph. This was achieved using a custom Matlab based fitting routine that uses a Gaussian fit centred on the active region to provide a peak intensity (see methods and (9)). The next step was to convert this intensity into the number of S1-GFP molecules bound. To do this, for one kymograph we generated four independent Markov chains using the RJMCMC algorithm, and from this we calculated the average mean pixel intensity values associated with that number of binders. This value depends on the maximum number fluorescence states (components) used in the RJMCMC algorithm. Since the fluorescence intensity is linear with the number of binders, this was used as a prior constraint for the RJMCMC algorithm.

The average of four simulated RJMCMC chains with two different numbers of components is shown in Table 1. For 7 components (equivalent to background plus 6 binders) and 9 components (equivalent to background plus 8 binders) clear linear increments between binders are seen (Figure S1). However, the 9-component simulation provides a more saturated Gaussian mixtures model. Further confirmation of the choice of component number derives from the weighting or relative abundance of these binders. For 9 components the predominant population is between 1 and 5 molecules per bound cluster but there is still a 17% chance for 6 to 8 binders (8% weighting for 6 binders, 5% for 7 binders and 4% for 8 binders). Therefore, the values from the 9-component simulation were used to define the fluorescence intensities for subsequent analysis.

**Table 1:** The mean, variance and weight of pixel intensity for a maximum of 6 or 8 myosin binders

|  | Total of 6 Binders (7 components) |  |  | Total of 8 Binders (9 components) |  |  |
| --- | --- | --- | --- | --- | --- | --- |
| | Mean=$\mu_6$ | Variance=$\sigma_6$ | Weight=$w_6$ | Mean=$\mu_8$ | Variance=$\sigma_8$ | Weight=$w_8$ |
| **0 Binders** | 57.17 | 34.73 | 0.12 | 24.52 | 35.58 | 0.10 |
| **1 Binder** | 150.75 | 42.07 | 0.21 | 138.57 | 41.54 | 0.15 |
| **2 Binders** | 198.26 | 50.68 | 0.20 | 173.87 | 46.04 | 0.16 |
| **3 Binders** | 254.73 | 60.71 | 0.15 | 210.90 | 51.35 | 0.16 |
| **4 Binders** | 314.23 | 69.58 | 0.14 | 264.22 | 55.51 | 0.14 |
| **5 Binders** | 401.37 | 85.73 | 0.11 | 322.20 | 62.04 | 0.11 |
| **6 Binders** | 551.86 | 111.40 | 0.07 | 382.67 | 80.58 | 0.08 |
| **7 Binders** | - | - | - | 490.73 | 72.01 | 0.05 |
| **8 Binders** | - | - | - | 652.38 | 91.13 | 0.04 |

From the values in Table 1 we take the simplest approach for choosing the cut-off pixel intensity value of each population of binders by using the midpoint between successive means for each of the 9 components. For example, the cut-off values for pixel intensity from baseline to binder 1 (values in table 1: 24.52 to 138.57) is 81.55 and between binders 1 to 2 (values in table 1: 138.57 to 173.87) is 156.21; therefore any peaks in the kymograph within this intensity range would be assigned as a single bound myosin. The mean (Table 1) and cut-off values (Table S1) were used for analysis of all subsequent kymographs to convert the intensity values of fitted peaks in the data to the number of bound myosins.

### Probabilities of transitioning from one state to another

The purpose of this analysis was to understand the probability of S1-GFP leaving from or associating with an active region. Knowing the number of myosins in each active region at every point in time enabled us to calculate transition matrices, which are tables that indicate what happens when the occupancy of an active region changes. To generate a transition matrix we examined the fate of each active region or cluster by measuring the frequency of transitions from one cluster size to another within a single frame (vertical slice of the kymograph). From these measurements we generated a matrix of probabilities (Figure 3a). The rows represent the final cluster size and the columns the starting size. The central diagonal is zero since we are measuring movement away from the current cluster size. All rows sum to one and there are no vertical transitions. For example, starting with a cluster size of four molecules in Figure 3a, the probability of releasing an S1-GFP is 14.6% whereas the probability of a single S1-GFP binding is 5.5%. It can be seen that near the central diagonal there is an increased probability, corresponding to the release/binding of single S1-GFPs. However, larger transition probabilities are seen in the first column, indicating a high propensity for complete detachment of all molecules in a single frame. By contrast, S1-GFP binding does not appear concerted, since there is very little probability of forming complexes in the top right of the matrix.

**Figure 3.**
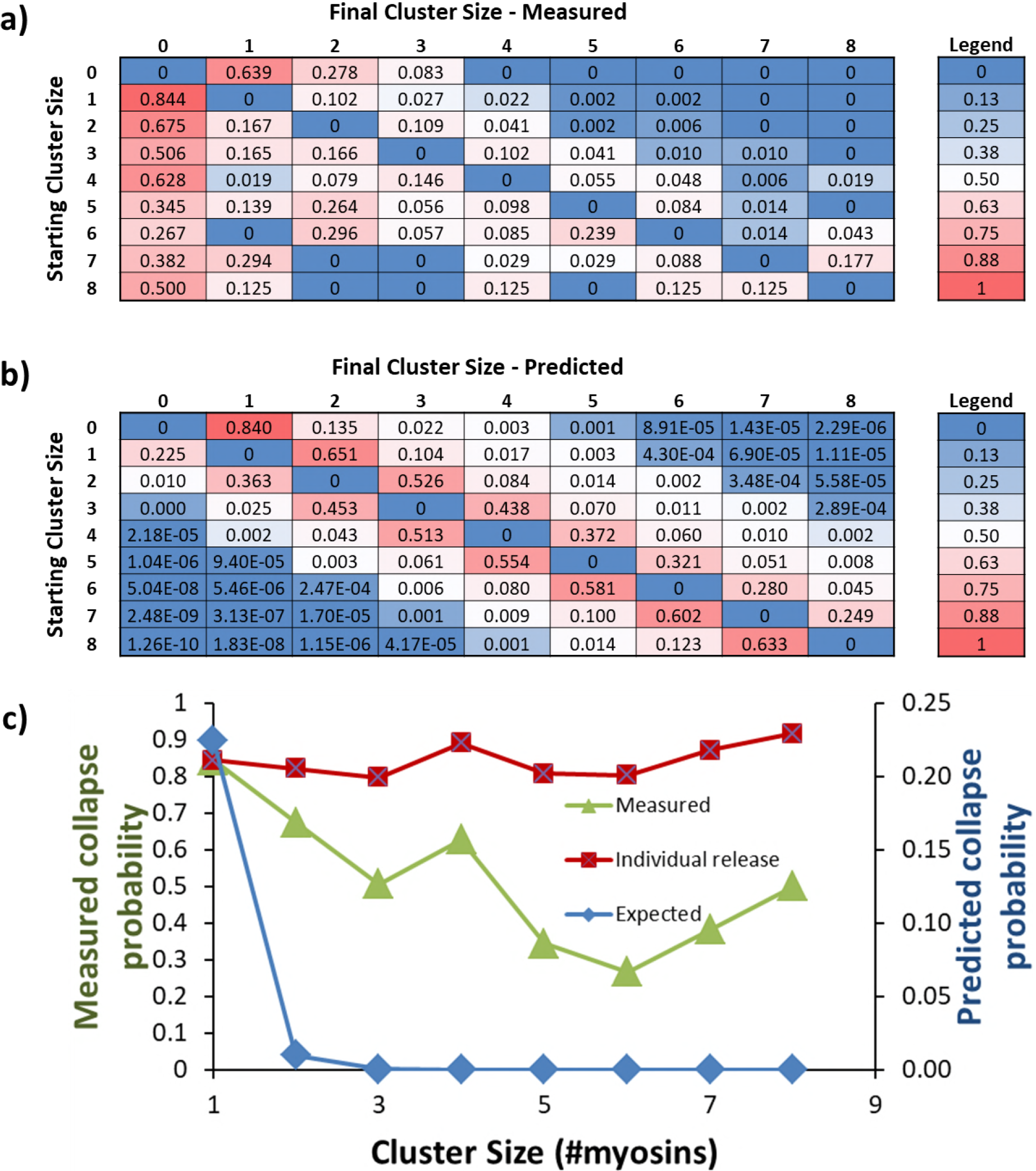
Predicted and measured transitions between cluster sizes. Using an RJMCMC model to determine the number of myosins in each cluster the transition rates between them can be determined. ***a***) From these rates a transition probability table is constructed where the rows are starting cluster size and the columns final. The central diagonal is zero because this table measures the probability of leaving that cluster, all numbers to the right of the diagonal are binding events and to the left are detachments. ***b***) Using the probability of association empirically determined from (a) and known kinetic parameters that govern myosin release it is possible to construct an expected table of transition probabilities (see discussion for more details). The overall pattern of behaviour between measured (a) and predicted (b) is similar with the exception of the first column, which indicates that the probability of release of all myosins is much greater than predicted. ***c***) A graph of probability of collapse versus initial cluster size. The predicted values from the first column of (b) show the expected power law dependence correlates with cluster size (blue). For the measured data (green) there is considerable deviation indicating a much higher probability of collapse, decreasing with larger clusters. These values can be corrected by the cluster size (see discussion) to yield the probability of individual myosin release (red, values associated with left axis). These are not seen to substantially change, indicating the myosin release rate is accelerated identically regardless of cluster size.

## Discussion

Classical cooperative systems use the energy of substrate binding to activate association of further substrate, all within the confines of the structural unit. These confines determine the number of substrate molecules that can bind (25–27). The sarcomeric thin filament presents a uniquely cooperative environment, where the number of substrate molecules, in this case myosin, that can associate is not structurally defined. Furthermore, this system is allosterically controlled through calcium binding to troponin which, together with tropomyosin, controls myosin′s access to binding sites on actin (2). Myosin binding results in a more classical cooperative activation as it holds Tm away from adjacent sites thus propagating further myosin association (9, 28–31). Despite this understanding, many crucial details remain to be understood, including how the thin filament relaxes. This process occurs as the calcium concentration drops in the myocyte, leading to re-binding of troponin I and repression of myosin binding (32). Given the very high local concentrations of myosin in the sarcomere (upwards of 0.15 mM (13)), troponin I binding and Tm movement will compete with myosin re-binding, making relaxation difficult. To reveal this mechanism, we have studied the dynamic process of metastable myosin binding that occurs near the mid-point of the calcium activation curve. In these conditions we directly observe single molecules of fluorescently tagged myosin S1 interacting with the thin filament, forming clustered regions of activation. Detailed examination using a Reversible Jump Monte Carlo Markov Chain approach reveals an unexpectedly high probability of concerted myosin release, suggesting an active process of myosin detachment that we term catastrophic collapse.

### Myosin detachment from active regions is cooperative

Activation occurs through the initial association of myosin to open the thin filament for subsequent myosins to bind (9). These bind in a collision limited process and, although stochastic, the process of activation is predictable. Likewise, the release of myosin from the thin filament is predicted from the ADP release rate constant and the second order ATP binding rate constant. To determine the association probability empirically we used the information embedded in the transition matrix shown in Figure 3a. Because of the stochastic nature of binding we used the mean probability for the first binder (the first value to the right of the zero diagonal in Figure 3a) as an estimate of the association probability. The probability of association will scale according to the power of the number of attached molecules. Therefore, if two myosins bind the expected probability for this event is square that of a single binder, for three myosins it is the cubed probability etc. Using this analysis, it is possible to calculate the expected probability for any number of myosins binding in one movie frame to an existing active region, shown in Figure 3b as the values to the right of the diagonal. The probability of detachment is calculated from the parameters that govern attachment time, ADP release and ATP binding. ADP release for skeletal muscle myosin is extremely fast by comparison with ATP binding at 0.1 μM ATP; therefore, we only calculated the probability of detachment from the latter. Using a second order ATP binding rate constant of 1.9 μMs^−1^ (9, 10), the detachment rate constant is calculated as 0.19 s^−1^. The duration of each frame is 300 ms therefore the expected probability of detachment during a single frame is (1-e^(−0.19*0.300)^) = 5.6%. As with attachment, this value will scale according to the number of myosins in a cluster; however, any molecule can leave the cluster, therefore the probability of detachment becomes the product of a single detachment and the active region size, scaled by a term which takes into account the possible combinations of myosins detaching from a cluster (see the supplementary information). Using these conditions, we created the rest of the table of transition probabilities in Figure 3b. The interaction between myosin and the thin filament is clear from this table; there is an increasingly lower probability of dis/association further away from the central diagonal, suggesting that in a stochastic model one would expect active regions to increase or decrease in size primarily in single steps. Our temporal resolution during imaging (3.3 fps) is sufficient to detect stepwise detachment based on the lifetime calculated above. However, when we calculated the observed transition rates using the RJMCMC model for our measured data (Figure 3a), the first column is strikingly different from the predicted behaviour (Figure 3b). The heat map in Figure 3a indicates that the most likely outcome in the conditions of metastable myosin binding is complete detachment of all myosins. This confirms the observed large decreases in the raw data intensity of the active region (Figure 2), indicating several myosins leaving simultaneously. Photobleaching cannot explain this synchronous reduction in fluorescence. Therefore these data indicate that myosin release from thin filaments is concerted.

### Tropomyosin accelerates the detachment of myosin to the same degree

The data presented here suggest that myosin′s detachment from actin is accelerated by Tm. To determine if this acceleration is uniform or if the active region size affects detachment we plotted the probability of catastrophic collapse versus initial cluster size for both the measured (green, Figure 3c) and predicted (blue, Figure 3c) data. There is a reduction in the probability of collapse with increasing initial cluster size (green, Figure 3c); however, to determine if individual myosins are more readily detached from larger clusters we recalculated the probability of release to account for the cluster size. If multiple myosins are releasing in a single step the probability of each myosin releasing is the n^th^ root of the collapse probability, where n is the active region cluster size. Across cluster sizes from 1 to 8 the probability of individual myosin detachment is approximately the same (red, Figure 3c), with a mean value of 84.4 ± 0.04 %. As calculated above we predict 0.19 s^−1^ for the detachment rate constant in these conditions, therefore to achieve an unbinding probability of 84 % the detachment rate constant would be accelerated to ˜6.1 s^−1^. Measurements of the detachment rate constant at low calcium concentrations (10) have not shown an increased detachment rate constant. However, since the slowest rate constant determined was approximately 5 s^−1^, similar to that observed here, it may be accelerated release would not be evident unless the [ATP] is dropped further. At saturating ATP concentrations the detachment of heads would be much faster than 6.1 s^−1^, however under load, lifetimes are reduced possibly reaching this threshold. The precise mechanism of how catastrophic collapse occurs on the thin filaments is not revealed in these experiments because of the limitations of wide-field imaging. Future studies using super-resolution techniques may reveal the location of the last myosin to be released within the active region; or may reveal that structural changes in actin assist in concerted release. Either way the observation of catastrophic collapse provides a new and important parameter for models of muscle contraction.

## Conclusion

In this study we have investigated the behaviour of myosin S1 molecules binding metastably to regulated thin filaments using a single molecule imaging approach, offering new insight into how the thin filament relaxes. From our results it is evident that active regions are turned off in a concerted fashion, a mechanism we term catastrophic collapse. The role of this ′catastrophic collapse′ in the process of muscle contraction remains to be determined, although this nuanced understanding of relaxation may help in our understanding of disease. Hypertrophic cardiomyopathy (HCM) is characterised by a deficit in relaxation (5, 33–36) that is even evident in the first images of muscle cross-sections where glycogen granules were noted to be absent (37). Mutations that reduced the stiffness of tropomyosin (38) could impair catastrophic collapse providing a molecular explanation for incomplete relaxation during HCM. In such cases the thin filament becomes less able to release the myosin heads and completely relax.

## Author contribution

A.V.I collected all data, performed data analysis and wrote the manuscript; M.M. and H.D. carried out the RJMCMC analysis and wrote the manuscript; N.M.K. conceived the experiments, analysed the data and wrote the manuscript.

## Acknowledgments

We thank Rama Desai for collecting preliminary data for this project and for advice. We also thank the Kad lab members for useful discussions. This project was funded by the British Heart Foundation (FS/13/69/30504). The authors declare no conflicts of interest.

## References

1. Poole,K. J., M. Lorenz, G. Evans, G. Rosenbaum, A. Pirani, R. Craig, L. S. Tobacman, W. Lehman, and K. C. Holmes. 2006. A comparison of muscle thin filament models obtained from electron microscopy reconstructions and low-angle X-ray fibre diagrams from nonoverlap muscle. J Struct Biol 155(2):273–284.

2. Gordon, A. M., E. Homsher, and M. Regnier. 2000. Regulation of contraction in striated muscle. Physiol Rev 80(2):853–924.

3. Linari, M., E. Brunello, M. Reconditi, L. Fusi, M. Caremani, T. Narayanan, G. Piazzesi, V. Lombardi, and M. Irving. 2015. Force generation by skeletal muscle is controlled by mechanosensing in myosin filaments. Nature 528(7581):276–279.

4. Reconditi, M., M. Caremani, F. Pinzauti, J. D. Powers, T. Narayanan, G. J. Stienen, M. Linari, V. Lombardi, and G. Piazzesi. 2017. Myosin filament activation in the heart is tuned to the mechanical task. Proc Natl Acad Sci U S A 114(12):3240–3245.

5. Tardiff, J. C. 2005. Sarcomeric proteins and familial hypertrophic cardiomyopathy: linking mutations in structural proteins to complex cardiovascular phenotypes. Heart Fail Rev 10(3):237–248.

6. McKillop, D. F., and M. A. Geeves. 1993. Regulation of the interaction between actin and myosin subfragment 1: evidence for three states of the thin filament. Biophys J 65(2):693–701.

7. Walcott, S., and N. M. Kad. 2015. Direct Measurements of Local Coupling between Myosin Molecules Are Consistent with a Model of Muscle Activation. PLoS Comput Biol 11(11):e1004599.

8. Maytum, R., S. S. Lehrer, and M. A. Geeves. 1999. Cooperativity and switching within the three-state model of muscle regulation. Biochemistry 38(3):1102–1110.

9. Desai, R., M. A. Geeves, and N. M. Kad. 2015. Using fluorescent myosin to directly visualize cooperative activation of thin filaments. J Biol Chem 290(4):1915–1925.

10. Geeves, M. A., and S. S. Lehrer. 1994. Dynamics of the muscle thin filament regulatory switch: the size of the cooperative unit. Biophys J 67(1):273–282.

11. Heeley, D. H., B. Belknap, and H. D. White. 2002. Mechanism of regulation of phosphate dissociation from actomyosin-ADP-Pi by thin filament proteins. Proc Natl Acad Sci U S A 99(26):16731–16736.

12. Heeley, D. H., B. Belknap, and H. D. White. 2006. Maximal activation of skeletal muscle thin filaments requires both rigor myosin S1 and calcium. J Biol Chem 281(1):668–676.

13. Ferenczi, M. A., E. Homsher, and D. R. Trentham. 1984. The kinetics of magnesium adenosine triphosphate cleavage in skinned muscle fibres of the rabbit. J Physiol 352(1):575–599.

14. Pardee, J. D., and J. A. Spudich. 1982. Purification of muscle actin. Methods Enzymol 85 Pt B:164–181.

15. Monteiro, P. B., R. C. Lataro, J. A. Ferro, and F. D. C. Reinach. 1994. Functional Alpha-Tropomyosin Produced in Escherichia-Coli-a Dipeptide Extension Can Substitute the Amino-Terminal Acetyl Group. J Biol Chem 269(14):10461–10466.

16. Janco, M., A. Kalyva, B. Scellini, N. Piroddi, C. Tesi, C. Poggesi, and M. A. Geeves. 2012. alpha-Tropomyosin with a D175N or E180G mutation in only one chain differs from tropomyosin with mutations in both chains. Biochemistry 51(49):9880–9890.

17. Homsher, E., B. Kim, A. Bobkova, and L. S. Tobacman. 1996. Calcium regulation of thin filament movement in an in vitro motility assay. Biophys J 70(4):1881–1892.

18. Margossian, S. S., and S. Lowey. 1982. Preparation of myosin and its subfragments from rabbit skeletal muscle. Methods Enzymol 85 Pt B:55–71.

19. Springall, L., A. V. Inchingolo, and N. M. Kad. 2016. DNA-Protein Interactions Studied Directly Using Single Molecule Fluorescence Imaging of Quantum Dot Tagged Proteins Moving on DNA Tightropes. Methods Mol Biol 1431:141–150.

20. Kad, N. M., H. Wang, G. G. Kennedy, D. M. Warshaw, and B. Van Houten. 2010. Collaborative dynamic DNA scanning by nucleotide excision repair proteins investigated by singlemolecule imaging of quantum-dot-labeled proteins. Mol Cell 37(5):702–713.

21. Tokunaga, M., N. Imamoto, and K. Sakata-Sogawa. 2008. Highly inclined thin illumination enables clear single-molecule imaging in cells. Nat Methods 5(2):159–161.

22. Konopka, C. A., and S. Y. Bednarek. 2008. Variable-angle epifluorescence microscopy: a new way to look at protein dynamics in the plant cell cortex. Plant J 53(1):186–196.

23. Feng, D. I., 2016. Package ‘miscF’. https://www.rdocumentation.org/packages/miscF

24. Plummer, M., N. Best, K. Cowles, and K. Vines. 2010. coda: Output analysis and diagnostics for MCMC. R package version 0.14–2.

25. Monod, J., J. Wyman, and J. P. Changeux. 1965. On the Nature of Allosteric Transitions: A Plausible Model. J Mol Biol 12:88–118.

26. Koshland, D. E., Jr., G. Nemethy, and D. Filmer. 1966. Comparison of experimental binding data and theoretical models in proteins containing subunits. Biochemistry 5(1):365–385.

27. Cliff, M. J., N. M. Kad, N. Hay, P. A. Lund, M. R. Webb, S. G. Burston, and A. R. Clarke. 1999. A kinetic analysis of the nucleotide-induced allosteric transitions of GroEL. J Mol Biol 293(3):667–684.

28. Craig, R., and W. Lehman. 2001. Crossbridge and tropomyosin positions observed in native, interacting thick and thin filaments. J Mol Biol 311(5):1027–1036.

29. Trybus, K. M., and E. W. Taylor. 1980. Kinetic studies of the cooperative binding of subfragment 1 to regulated actin. Proc Natl Acad Sci U S A 77(12):7209–7213.

30. Swartz, D. R., and R. L. Moss. 1992. Influence of a strong-binding myosin analogue on calcium-sensitive mechanical properties of skinned skeletal muscle fibers. J Biol Chem 267(28):20497–20506.

31. Kad, N. M., S. Kim, D. M. Warshaw, P. VanBuren, and J. E. Baker. 2005. Single-myosin crossbridge interactions with actin filaments regulated by troponin-tropomyosin. Proc Natl Acad Sci U S A 102(47):16990–16995.

32. Galinska-Rakoczy, A., P. Engel, C. Xu, H. Jung, R. Craig, L. S. Tobacman, and W. Lehman. 2008. Structural basis for the regulation of muscle contraction by troponin and tropomyosin. J Mol Biol 379(5):929–935.

33. Bai, F., A. Weis, A. K. Takeda, P. B. Chase, and M. Kawai. 2011. Enhanced active cross-bridges during diastole: molecular pathogenesis of tropomyosin’s HCM mutations. Biophys J 100(4):1014–1023.

34. Moore, J. R., L. Leinwand, and D. M. Warshaw. 2012. Understanding cardiomyopathy phenotypes based on the functional impact of mutations in the myosin motor. Circ Res 111(3):375–385.

35. Sewanan, L. R., J. R. Moore, W. Lehman, and S. G. Campbell. 2016. Predicting Effects of Tropomyosin Mutations on Cardiac Muscle Contraction through Myofilament Modeling. Front Physiol 7(473):473.

36. Sen-Chowdhry, S., D. Jacoby, J. C. Moon, and W. J. McKenna. 2016. Update on hypertrophic cardiomyopathy and a guide to the guidelines. Nat Rev Cardiol 13(11):651–675.

37. Teare, D. 1958. Asymmetrical hypertrophy of the heart in young adults. Br Heart J 20(1):1–8.

38. Li, X. E., W. Suphamungmee, M. Janco, M. A. Geeves, S. B. Marston, S. Fischer, and W. Lehman. 2012. The flexibility of two tropomyosin mutants, D175N and E180G, that cause hypertrophic cardiomyopathy. Biochem Biophys Res Commun 424(3):493–496.

